# A voltage-dependent depolarization induced by low external glucose in neurons of the nucleus of the tractus solitarius of rats: interaction with K_ATP_ channels regulated by external glucose

**DOI:** 10.1101/328658

**Authors:** Cahuê de Bernardis Murat, Ricardo Mauricio Leão

**Author notes:** Corresponding author: Departamento de Fisiologia. FMRP-USP. Av. Bandeirantes 3900. Ribeirão Preto – SP. 14049-900. Brazil.

## Abstract

The brainstem nucleus of the tractus solitarius (NTS) is an integrative center for autonomic counterregulatory responses to hypoglycemia, NTS neurons can also sense fluctuations in extracellular glucose levels altering their membrane potential. K_ATP_ channels links the metabolic status of the neuron to its excitability, but the role of K_ATP_ channels in controlling NTS neurons excitability and in sensing extracellular glucose changes is not clear. Here we investigated using *in vitro* electrophysiological recordings in brainstem slices the influence of K_ATP_ channels on the membrane potential of NTS neurons in normoglicemic and hyperglycemic external glucose concentrations, and after switching the external glucose to a hypoglycemic level. We found that in normoglicemic (5 mM) external glucose application of tolbutamide, a K_ATP_ blocker, induced a substantial depolarization of most NTS neurons, while application of diazoxide, a K_ATP_ opener, hyperpolarized the membrane of all NTS neurons. Interestingly, neurons not responsive to tolbutamide were more depolarized than responsive neurons. In a hyperglycemic solution (10 mM glucose) few neurons depolarized in response to tolbutamide. We found that these neurons were more depolarized than neurons in 5 mM glucose and only the more hyperpolarized responded to tolbutamide. The non-responsive neurons did not respond to tolbutamide even when hyperpolarized. Interestingly application of a low-gucose solution (0.5 mM) did not hyperpolarized the RMP but produced a depolarization in most neurons. This effect was voltage-dependent not seen in neurons more depolarized, but could be observed when the neurons were hyperpolarized. Depolarization by tolbutamide avoided further depolarization by low glucose, unless the membrane was hyperpolarized. Application of 0.5 mM glucose solution in neurons incubated in 10 mM glucose depolarized the membrane only in more hyperpolarized neurons, which responded to tolbutamide, or after membrane hyperpolarization. The effect of glucose was caused by activation of a cationic current with a reversal potential around the potential were the neurons were non-responsive to low glucose. We conclude that NTS neurons present K_ATP_ channels open at rest in normoglicemic conditions, and that their state is affected by extracellular glucose. Moreover, NTS neurons depolarize the membrane in response to the application of a low-glucose solution, but this effect is occluded by membrane depolarization triggered by K_ATP_ blockage. This suggests a homeostatic regulation of the membrane potential by glucose and a possible mechanism related to hypoglycemia-associated autonomic failure.

## Introduction

The brainstem of the nucleus of the tractus solitarius (NTS) is the primary central site for viscerosensory afferent fibers arising from peripheral neurons, including peripheral chemo-sensing neurons (Accorsi-Mendonça et al., 2009; Donovan & Watts, 2014). NTS neurons densely project to the dorsal motor nucleus of the vagus, where the cell bodies of the preganglionar parasympathetic branch of the vagus are located (Marty *et al.*, 2007; Verberne *et al.*, 2014). Local nutrient and metabolic signals (including glucose and metabolites) in the NTS, as well as direct modulation by receptor agonists/antagonists in NTS neurons, induce physiological responses for the regulation of blood glucose concentration (Ritter *et al.*, 2000; Lam *et al.*, 2010; Zhao *et al.*, 2012). The NTS is associated with the physiological responses to hypoglycemia, the counter-regulatory responses, which result in glucagon secretion, increased food intake behavior and increase in the sympathetic tone. For instance, hypoglycemia induces c-Fos immunoreactivity in NTS neurons, and portal-mesenteric deafferentation suppressed this effect (Bohland *et al.*, 2014). Inhibitors of glucose metabolism injected in the NTS also increased glucagon secretion and food intake (Ritter et al., 2000; Andrew et al., 2007). Additionally, low glucose increases firing activity in carotid sinus nerve (Gao *et al.*, 2014), which mostly propagates signals into the NTS. Therefore, NTS neurons integrate a variety of synaptic inputs and mediate a plethora of autonomic counter regulatory mechanisms in response to food ingestion and signals related to the glycemic state to ensure adequate levels of glucose, including feeding behavior, gastric motility, and hormonal secretion (Marty *et al.*, 2007; Donovan & Watts, 2014; Hermann *et al.*, 2014).

The NTS is an interesting region for detecting changes in extracellular glucose due to presence of fenestrated capillaries (Gross et al., 1990), and its proximity to the area postrema, a circumventricular organ. Several reports have shown that NTS neurons directly detect fluctuations in glucose levels in the extracellular milieu. For instance, Mimee and Fergunson (2015) showed that slightly more than half of NTS neurons change their resting membrane potential to both increase and decrease in external glucose *in vitro*, and can depolarize or hyperpolarize by increased glucose. On the other hand, Balfour et al., (2006) observed that only a minority of the NTS neurons changed membrane potential in response to a zero glucose solution. Lamy et al, (2014) showed a depolarization of the membrane potential in response to 0.5 mM external glucose, only in GABAergic neurons in the NTS. Finally, McDougal et al., (2013) showed that a fraction of glia and neurons of NTS increased intracellular calcium in response to low glucose and inhibitors of glycolysis.

ATP-sensitive potassium (K_ATP_) channel links metabolic status and electrical excitability (Nichols, 2006), as classical described in pancreatic beta-cells (Ashcroft & Rorsman, 2013). This channel is a hetero-octameric complex comprised of four pore-forming K_ir_6.x subunits and four regulatory sulfonylurea receptor (SURx) subunits, and it conducts an inwardly rectifying potassium current that is inhibited by ATP binding to K_ir_6.x subunits and stimulated by ADP interaction with nucleotide-binding sites within SURx subunits (Hibino *et al.*, 2010). In the NTS, the subunits K_ir_6.2 and SUR1 are expressed in glucose-sensing neurons (Balfour *et al.*, 2006; Halmos *et al.*, 2015). Glucose-excited NTS neurons indeed appear to respond to changes in glucose levels using a mechanism similar to pancreatic beta-cells, in a GLUT/glucokinase/ K_ATP_ channels system (Thorens, 2012; Ashcroft & Rorsman, 2013), since K_ATP_ channel antagonists blunts the responsive of these neurons to increased glucose levels (Balfour *et al.*, 2006; Boychuk *et al.*, 2015).

Here we aimed to investigate how the membrane potential of NTS neuros is affected by changing glucose concentrations and the role of K_ATP_ channels. We found that NTS neurons depolarize in response to low-external glucose only when their membrane potential is hyperpolarized below −50 mV. Incubation in high glucose external solution depolarize the neurons by blocking K_ATP_ channels, and blunts their response to low-glucose. This unexpected finding show that NTS neurons in the presence of hyperglycemic glucose concentrations do not respond to fast reductions in external glucose, with implications to situations of prolonged hyperglycemia as diabetes mellitus.

## Materials and Methods

### Brainstem slices preparation

Animal procedures were performed according to protocol approved by the Committee on Ethics in Animal Experimentation (CEUA) from the School of Medicine of Ribeirão Preto, University of São Paulo (protocol # 149/2015). Brainstem slices containing the NTS were obtained as previously described (Accorsi-Mendonça et al., 2009). Male Wistar rats (3-to 11-week-old) were decapitated following isoflurane anesthesia, and the brainstem quickly removed and placed in a dish containing ice-cold artificial cerebrospinal fluid (aCSF) modified for slicing, containing (in mM): 87 NaCl, 2.5 KCl, 1.25 NaH_2_PO_4_, 25 NaHCO_3_, 75 sucrose, 25 D-glucose, 0.2 CaCl_2_, and 7 MgCl_2_ [330 mOsm/kg H_2_O, pH 7.4 when bubbled with carbogenic mixture (95% O_2_ and 5% CO_2_)]. The specimen was glued to the sectioning stage, and submerged in ice-cold slicing aCSF, in a vibratome (Vibratome 1000 Plus, Vibratome) chamber. Then, three to four coronal brainstem slices (250 μM) containing the NTS near the level of the area postrema (i.e., ± 500 μm rostral and caudal) were sectioned and incubated at 32 – 33°C for 45 min in aCSF, containing (in mM): 125 NaCl, 2.5 KCl, 1.25 NaH_2_PO_4_, 25 NaHCO_3_, 5 D-glucose, 2 CaCl_2_, and 1 MgCl_2_ [298 mOsm/kg H_2_O, pH 7.35 when bubbled with carbogenic mixture (95% O_2_ and 5% CO_2_)]. In some experiments, the aCSF contained 10 mM glucose. After this period, the slices were stored into the same solution at room temperature until use for electrophysiological recordings. Low-glucose (0.5 mM) aCSF, as well as 10 mM glucose aCSF, were made using equimolar amounts of NaCl or sucrose to maintain osmolality equal of the 5 mM glucose aCSF. No differences were observed in both conditions.

### Electrophysiological recordings

Single brainstem slices were transferred to a chamber mounted on a stage of an upright microscope (BX51WI; Olympus, Japan), and continuously perfused with aCSF at 30 – 33°C using an inline heating system (TC-324B; Warner Instruments, Hamden, CT, USA) at a rate of ∼2 mL/min using a gravity-driven perfusion system. Neurons were then visualized under DIC optics with a 60 x immersion objective, and patched with eletctrodes made with thick-walled borosilicate glass (BF150-86-10; Sutter Instruments, Novato, CA, USA) pulled using a horizontal puller (P-87; Sutter Instruments). The electrodes were filled with an internal solution, containing (in mM): 128 K-gluconate, 8 KCl, 10 HEPES, 0.5 EGTA, 4 Mg_2_ATP, 0.3 Na_2_GTP, and 10 Na-phosphocreatine (295 mOsm/kg H_2_O, pH 7.3), resulting in pipette tip resistance between 3 and 7 MΩ.

Whole-cell patch-clamp recordings were performed with an EPC-10 patch-clamp amplifier (HEKA Eletronik, Lambrecht, Germany) using the PatchMaster acquisition software (HEKA Eletronik), in current-clamp mode. Data was acquired at 20 kHz and low-pass filtered at 5 kHz (Bessel). Cells with series resistance larger than 30 MΩ or showing large variations during whole-cell recording were discarded. After entering the whole-cell configuration, neurons were allowed to stabilize their membrane potential for ∼10 min. Membrane potential was monitored for 10 min in aCSF with 5 or 10 mM glucose prior to switch to low glucose (0.5 mM) solution, or with drug, for 10 min before returning to 5 or 10 mM glucose aCSF. In some experiments, we perfused the low-glucose solution for more prolonged periods (20-30 minutes). Hyperpolarizing pulses (ranging from −30 to −60 pA; 1000 ms) were applied every 15 seconds to monitor input resistance and membrane time constant. All conditions were recorded during 10 min, and we just recorded one neuron per slice. Based on previous reports regarding glucose sensing in NTS neurons (Balfour *et al.*, 2006; Lamy *et al.*, 2014; Boychuk *et al.*, 2015), and the response profile to a low-glucose challenge observed in the current investigation, we assumed that a neuron was responsive to low glucose when they showed a clear membrane depolarization induced by low-glucose solution and/or drug. Based on our observations, we considered responsive neurons that presented a depolarization of at least 3 mV.

For voltage-clamp experiments, neurons were clamped at −70 mV and the membrane potential varied from −125 mV to −65 mV in seven steps of 500 ms.

### Drugs

Tetrodotoxin (0.5 μM) was purchased from Alomone Labs (Jerusalen, Israel), and tolbutamide (100 μM), diazoxide (200 µM), picrotoxin (100 µM), DLAP5 (40 µM), strychnine (1 µM) and 6,7-dinitroquinoxaline-2,3-dione (DNQX; 10 µM) were purchased from Sigma (St. Louis, MO). All drugs were diluted at the time of the experiment from 1000 x concentrated stock solutions in DMSO or water.

### Data Analysis

Electrophysiological data were analyzed with IGOR Pro 6.37 (WaveMetrics, Portland, OR, USA), MiniAnalysis 6.0 (Synaptosoft, Fort Lee, NJ, USA), and ClampFit 10.6 (Molecular Devices, Sunnyvale, CA, USA). Voltages were corrected for measured liquid junction potential of 10 mV. Data are showed as mean ± SEM. All data values were determined after obtained a plateau response. Resting membrane potential was analyzed as the mean of all points histogram of the recording taken in a segment of the last 3 - 4 min. Statistical analyses were performed using GraphPad Prism 6.0 (GraphPad Software, La Jolla, CA, USA), conducted with paired and unpaired two-tailed t-test, and one-way repeated measures ANOVA with Fisher’s LSD test. Correlations were determined using a linear regression. Significance level was set at p < 0.05.

## Results

### NTS neurons incubated in 5 mM glucose express partially closed K_ATP_ channels

K_ATP_ channels are traditionally associated to coupling the cell energy status and electrical activity, triggering depolarization in high ATP/ADP ratio, and hyperpolarization in low ATP/ADP ratio (Nichols, 2006; Hibino *et al.*, 2010). NTS neurons express K_ATP_ channels, which are more responsive to metabolic ATP than to ATP provided by the whole-cell pipette solution, even in intracellular ATP concentrations of 3-4 mM, like we used in our recordings (Balfour *et al.*, 2006). In order to know if the K_ATP_ channels in the NTS neurons were active in neurons incubated in 5 mM external glucose, a near normoglycemic concentration, we applied their antagonist tolbutamide (100 µM; n = 15, seven animals) and measured the change in RMP. We found that the neurons were viable in aCSF with 5 mM glucose during the whole incubation period of the experiment (up to 5 hours).

We verified that in 10 neurons (67%) incubated with aCSF containing 5 mM of glucose, tolbutamide triggered a strong and fast depolarization (16.7 ± 2.7 mV, from a mean of −76.1 ± 2.6 mV to −59.4 ± 1.2 mV; p = 0.0002; **Figures 1A*i*** and **1A***ii*), accompanied with robust increased R_input_ (190.5 ± 55.2 MΩ, from a mean of 406.9 ± 79.2 MΩ to 597.4 ± 73.4 MΩ; p = 0.007; **Figure 1A*iii***), in accordance to its effect in blocking K_ATP_ channels. Additionally, five neurons (33%) did not change their RMP in response to tolbutamide (1.0 ± 0.5 mV, from a mean of −65.1 ± 2.4 mV to −64.1 ± 2.2 mV; p = 0.1; **Figures 1B*i*** and **1B***ii*) as well as membrane input resistance (R_input_) (14.8 ± 11.0 MΩ, from a mean of 525.8 ± 121.8 MΩ to 540.6 ± 129.1 MΩ; p = 0.2; **Figure 1B*iii***). Interestingly, we verified that neurons unresponsive to tolbutamide were significantly more depolarized than neurons responsive to tolbutamide (−65.2 ± 2.4 mV vs. −76.1 ± 2.5 mV, respectively; p = 0.019; **Figure 1C**). Accordingly, we found a negative correlation of RMP and tolbutamide depolarization (r^2^ = 0.79; p < 0.0001; **Figure 1D**). On the other hand, no correlation between the change of R_input_ induced by tolbutamide (ΔR_input_) and initial R_input_ (r^2^ = 0.2; p = 0.19. **Figure 1E**) was observed. Therefore, we conclude that NTS neurons present active K_ATP_ channels, which can control RMP. Because tolbutamide-insensitive neurons had more depolarized RMP, it is possible that K_ATP_ channels are already closed by endogenous ATP in these neurons (see next section).

**Figure 1.**
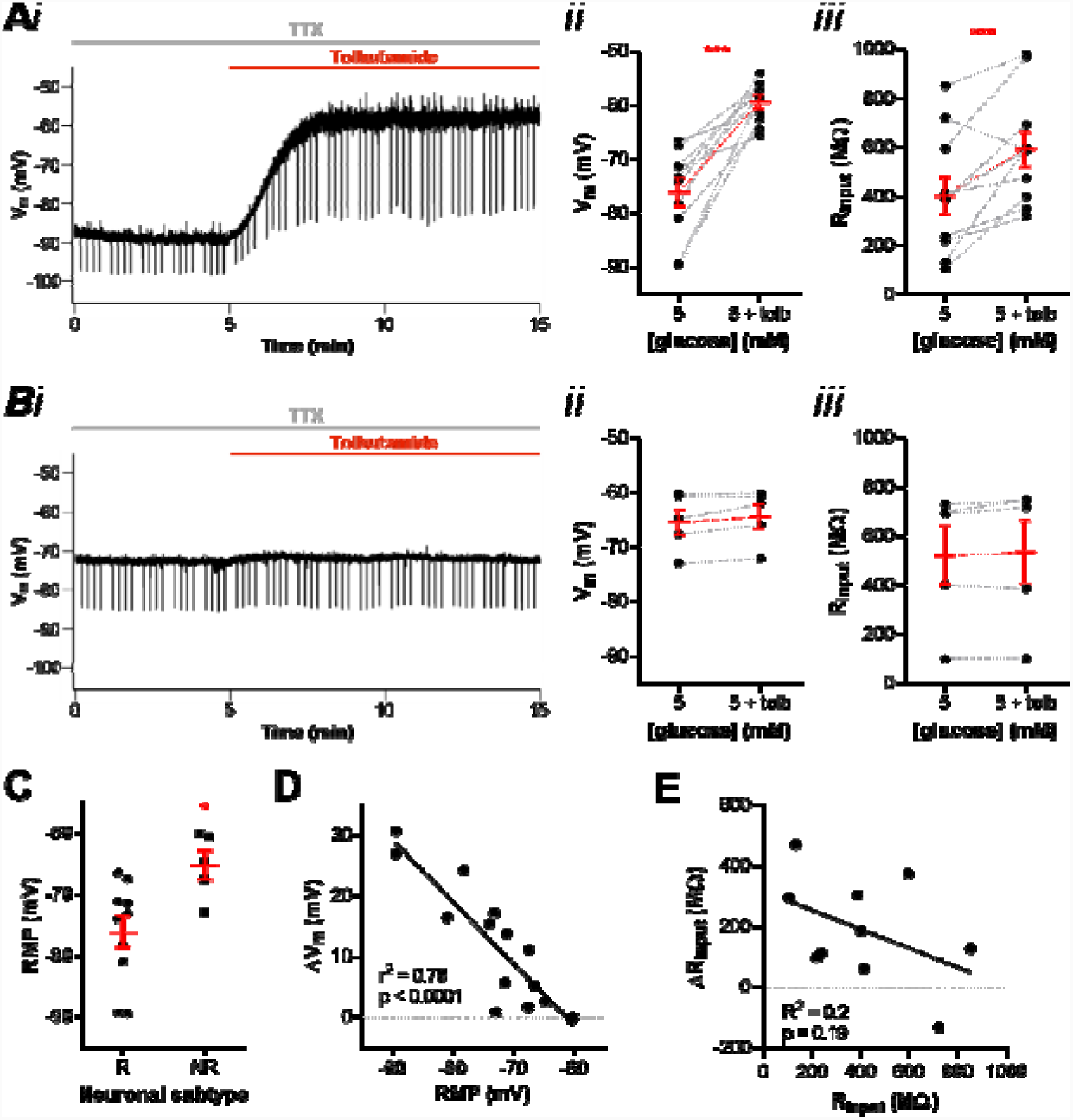
NTS neurons present active K_ATP_ channels. **A,** subset of neurons responsive to tolbutamide, as shown by a representative recording (*i*). The graphs show the summary of the tolbutamide effect on (*ii*) membrane potential (V_m_) and (*iii*) input resistance (R_input_) of neurons. **B,** subset of neurons unresponsive to tolbutamide, as shown by a representative recording (*i*). The graphs show the summary of the tolbutamide effect on (*ii*) V_m_ and (*iii*) R_input_ of neurons. **C,** comparison of the resting membrane potential (RMP) between neurons responsive (R) and unresponsive (NR) to tolbutamide. **D,** linear correlation between the change of V_m_ induced by tolbutamide and the RMP of neurons. **E,** linear correlation between the change of R_input_ induced by tolbutamide and the basal R_input_ of neurons. Tolb, tolbutamide; TTX, tetrodotoxin. *p < 0.05; ***p < 0.001.

To test if the non-responsive neurons express K_ATP_ channels we tested the effect of the K_ATP_ activator diazoxide (200 µM; n = 7, five animals) on the RMP of NTS neurons. Diazoxide produced a fast and partially reversible hyperpolarization of almost all NTS neurons tested (−8.8 ± 2.1 mV, from a mean of −73.7 ± 1.7 mV to - 82.6 ± 3.3 mV; p = 0.005; **Figure 2A*i***, **2A***ii*). Diazoxide also decreased R_input_ significantly (−230.4 ± 87.7 MΩ, from a mean of 532.3 ± 103.7 MΩ to 301.9 ± 65.3 MΩ; p = 0.04; Figure **2A***iii*) in accordance to the opening of an ion channel. Contrary to tolbutamide, we observed an effect of diazoxide in both depolarized and hyperpolarized NTS neurons. In fact, no correlation was observed between the effect of diazoxide on membrane potential and RMP (r^2^ = 0.24; p = 0.2; **Figure 2B**). However, we found a significant correlation between the change of R_input_ induced by diazoxide (ΔR_input_) and initial R_input_ (r^2^ = 0.61; p = 0.04; **Figure 2C**) suggesting that the magnitude of the effect of diazoxide is affected by the amount of K_ATP_ channels blocked. Thus, because diazoxide is able to open K_ATP_ blocked channels, we conclude that NTS neurons have K_ATP_ channels blocked at rest, and that modulation of K_ATP_ channels can affect RMP of NTS neurons bidirectionally.

**Figure 2.**
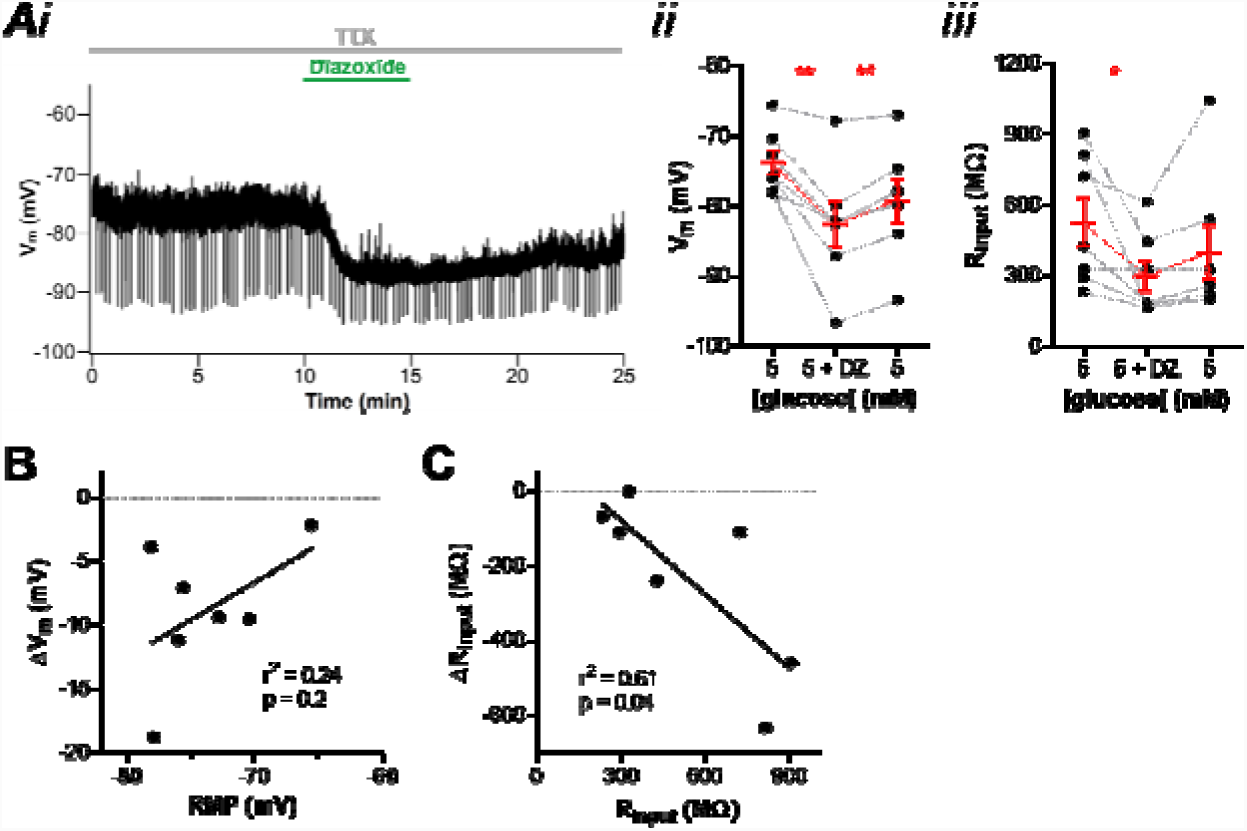
K_ATP_ channels control the resting membrane potential of NTS neurons. **A,** diazoxide induces a fast and partially reversible hyperpolarization of neurons, as shown by a representative recording (*i*). The graphs show the summary of the diazoxide effect on (*ii*) membrane potential (V_m_) and (*iii*) input resistance (R_input_) of neurons. **B,** linear correlation between the change of V_m_ induced by diazoxide and the resting membrane potential (RMP) of neurons. **C,** linear correlation between the change of R_input_ induced by diazoxide and the basal R_input_ of neurons. DZ, diazoxide; TTX, tetrodotoxin. *p < 0.05; **p < 0.01.

### Most NTS neurons in 5 mM external glucose depolarize in response to low-glucose aCSF

NTS neurons are involved in the counteregulatory response to hypoglycemia (Lamy et al., 2014). We showed that partially blocked K_ATP_ channels are present in NTS neurons, thus they can be subjected to modulation by the metabolic ATP/ADP ratio. Other groups showed that a fraction of NTS neurons changed membrane potential in response to changes in extracellular glucose. The effect of diazoxide suggests that there are K_ATP_ channels blocked in our basal conditions. We then tested if perfusion of a low external glucose solution could produce a hyperpolarization of NTS neurons by activation of K_ATP_ channels, caused by a reduction of the ATP/ADP levels. We then perfused the NTS neurons incubated in 5 mM glucose with a solution with low glucose (0.5 mM), a concentration that can be achieved in the CSF during periods of hypoglycemia (Seaquist et al;, 2001) and recorded the membrane potential. Surprisingly we found that perfusion of NTS neurons (n = 37, 23 animals) with a low-glucose (0.5 mM) solution depolarized most of NTS neurons (30 neurons; 81%). In these neurons the RMP was depolarized after perfusion of low glucose aCSF by 9.3 ± 1.0 mV, from an average of −74.3 ± 1.6 mV to −65.0 ± 1.7 mV (p < 0.0001; **Figure 3A*i*** and **3A***ii*). This effect took on average 383 ± 18 s to reach its peak, being reversible in most neurons (70%), and the RMP after returning to 5 mM glucose was - 71.3 ± 1.9 mV (p < 0.0001).

**Figure 3.**
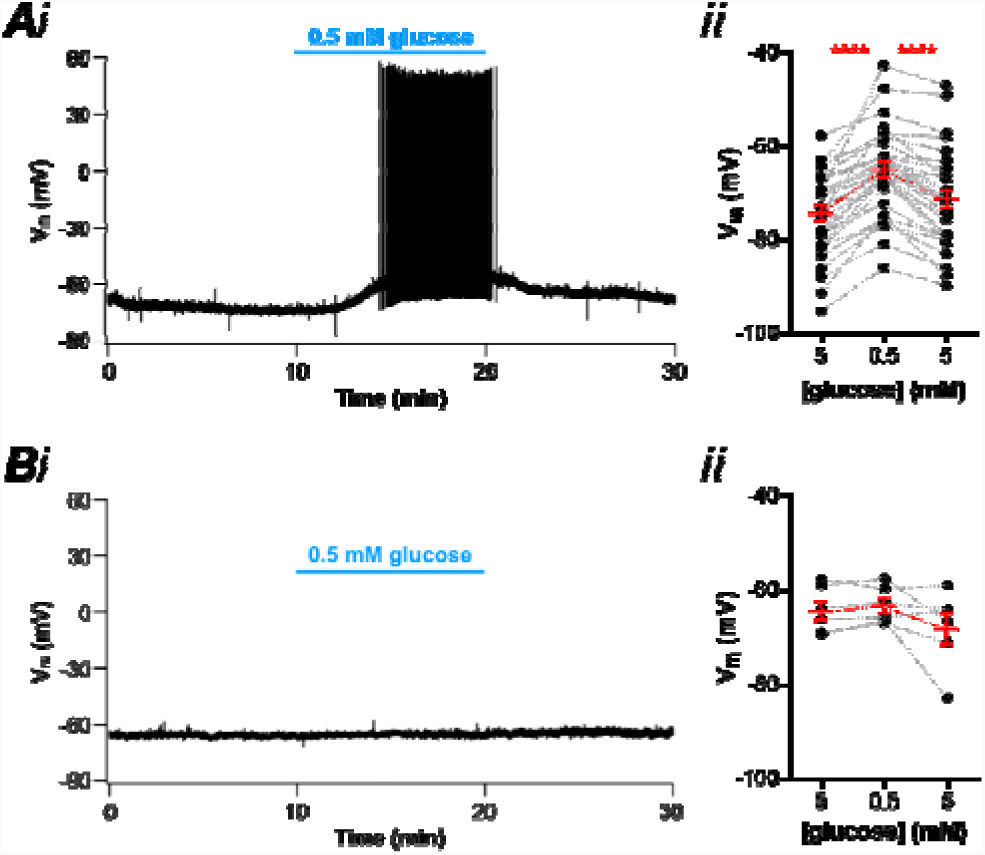
Most NTS neurons depolarize in response to a low-glucose challenge. **A,** subset of neurons depolarized by low glucose. **A***i*, representative recording of a responsive neuron showing a reversible depolarization and increased firing activity induced by low glucose. **A***ii*, summary of the low-glucose effect on the membrane potential (V_m_) of neurons. **B,** subset of neurons unresponsive to low glucose, as shown by a representative recording (*i*). **B***ii*, summary of the low-glucose effect on the V_m_ of neurons. ****p < 0.0001.

Depolarization induced by low glucose triggered action potential (AP) firing in 16 of 28 silent neurons (57%; 0.6 ± 0.2 Hz; p = 0.007), and an increase in AP frequency in the only two spontaneously active neurons (from a mean of 3.3 ± 2.7 Hz to 8.4 ± 2.0 Hz in low glucose). Surprisingly, only a single cell (3%) was hyperpolarized during perfusion with low-glucose solution, from −60 mV to −68 mV, and this effect was partially reversed (41%) on reinstatement of 5 mM external glucose. Finally, six neurons (16%), were considered unresponsive to low glucose with mean RMP in 5 mM glucose of −64.1 ± 2.0 mV, and after 0.5 mM glucose of - 63.1 ± 1.6 mV; a difference of 1.0 ± 0.7 mV; p = 0.2; **Figure 3B*i*** and **3B***ii*).

To test the dependence on activation of voltage-gated sodium channels for low-glucose sensing, we performed the low-glucose challenge in the presence of tetrodotoxin (TTX; 0.5 µM) in a second set of NTS neurons (n = 23 cells, 11 animals). Again, we identified three types of responses. Fourteen cells (61%) were depolarized after low glucose by 8.8 ± 1.0 mV, from a mean of −79.1 ± 1.6 mV to −70.3 ± 1.3 mV (p < 0.0001; **Figures 4A*i*** and **4A***ii*), and took, on average, 348 ± 18 s to peak. This effect was reversible in most neurons (71%), and the RMP after returning to 5 mM glucose was −77.0 ± 1.8 mV (p < 0.0001). A single cell (4%) was hyperpolarized by - 5.5 mV, from −72.1 mV to −77.6 mV in low glucose, and this effect was reverted after returning to 5 mM glucose. Lastly, eight cells (35%) were unresponsive to low glucose solution (1.1 ± 0.5 mV; from a RMP of −66.9 ± 1.6 mV to −65.8 ± 1.7 mV; p = 0.07; **Figures 4B*i*** and **4B***ii*). Both depolarizing effect and latency to peak response were not different from what was observed in experiments conducted with no TTX (p = 0.7, and p = 0.2, respectively), and as in normal aCSF, most neurons depolarized in low-glucose solution (84% in normal aCSF, and 65% in TTX). We conclude that the depolarization triggered by low glucose is not dependent on the activation of voltage-gated sodium channels and action potential firing. Most of the experiments shown from now on were performed in the presence of TTX.

**Figure 4.**
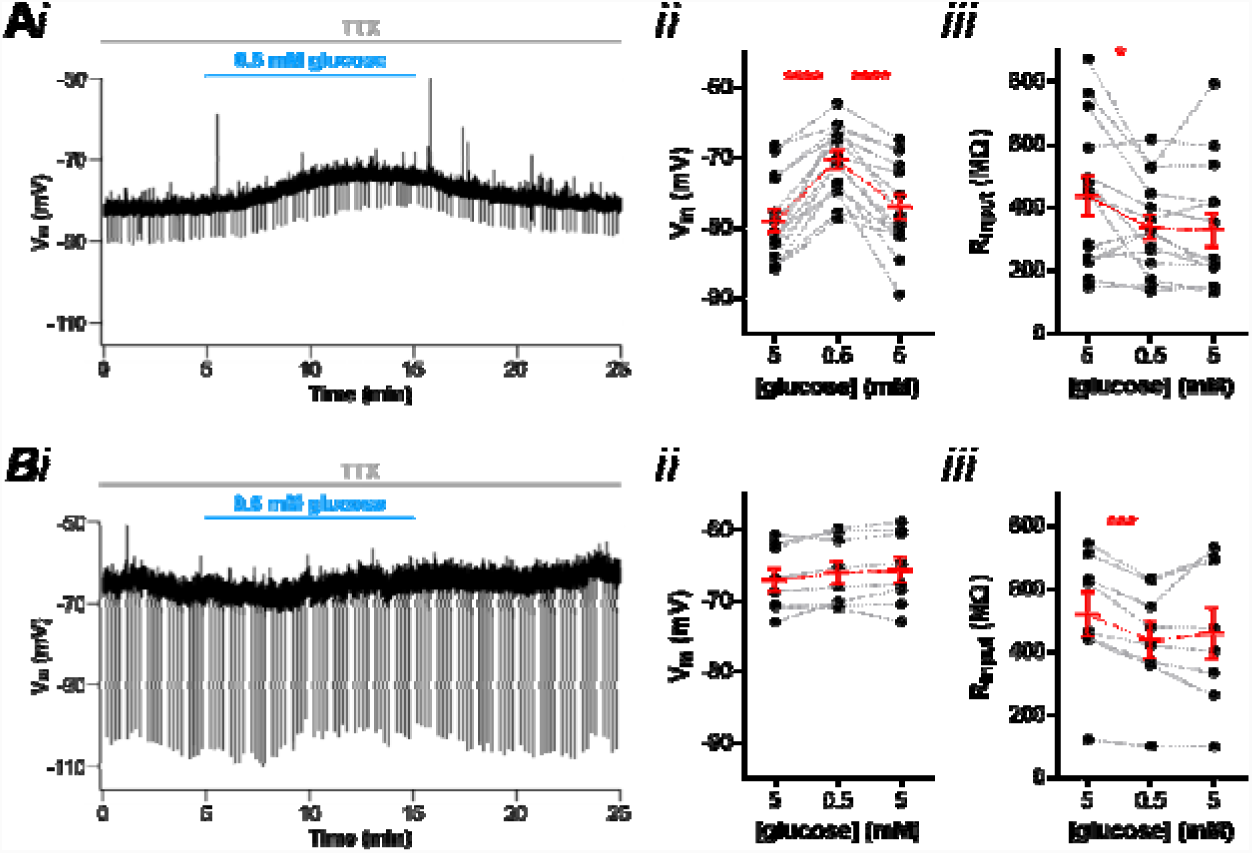
Low-glucose sensing of NTS neurons is independent of action potential-dependent neurotransmission. **A,** subset of neurons depolarized by low glucose in the presence of tetrodotoxin (TTX). **A***i*, representative recording of a responsive neuron showing a reversible depolarization and decreased input resistance (R_input_) induced by low glucose. The graphs show the summary of the low-glucose effect on (*ii*) membrane potential (V_m_) and (*iii*) R_input_ of neurons. **B,** subset of neurons unresponsive to low glucose in the presence of TTX, as shown by a representative recording (*i*). The graphs show the summary of the low-glucose effect on (*ii*) V_m_ and (*iii*) R_input_ of neurons. *p < 0.05; ***p < 0.001; ****p < 0.0001.

Lamy *et al*. (2014) found that the membrane depolarization induced by low glucose in GABAergic NTS neurons was accompanied by an increase in the membrane input resistance (R_input_) caused by inhibition of a potassium leak conductance. Contrary to the observation of Lamy et al., (2014) we verified that NTS neurons depolarized by low glucose showed a significant decrease in R_input_ (−101.1 ± 37.8 MΩ, from a mean of 440.3 ± 61.6 MΩ to 339.2 ± 40.0 MΩ; p = 0.02; **Figure 4A*iii***). Differently from the observed with the membrane potential, these effects did not revert on reinstatement of 5 mM glucose aCSF (p = 0.7). Additionally, neurons unresponsive to low glucose also showed a decrease in R_input_ (−80.3 ± 13.2 MΩ, from a mean of 522.9 ± 71.1 MΩ to 442.6 ± 61.6 MΩ; p = 0.0005; **Figure 4B*iii***). Like in responsive neurons, R_input_ did not revert after returning to 5 mM glucose aCSF in non-responsive cells (p = 0.5). Interestingly, the single neuron hyperpolarized by low-glucose in TTX showed a robust decrease in R_input_ (−629.2 MΩ, from 900.5 MΩ to 271.3 MΩ), suggesting the opening of K_ATP_ channels.

Interestingly, we observed that the depolarization caused by low glucose was strongly correlated with neuronal RMP (r^2^ = 0.53; p < 0.0001; **Figure 5A*i***), with the more negative the RMP, the greater the membrane potential change in response to low glucose. We then compared the RMP between responsive and non-responsive neurons, and found that the RMP of non-responsive cells were significantly more depolarized than responsive cells (−66.8 ± 1.6 mV vs. −78.6 ± 1.6 mV, respectively; p = 0.0001; **Figure 5B**). On the other hand, we found an inverse weak correlation of low glucose effect on ΔV_m_ and ΔR_input_ (r^2^ = 0.18, p = 0.04; **Figure 5A*ii***). We conclude that the depolarization induced by low external glucose uses a voltage-dependent mechanism, which might involve the opening of a depolarizing membrane conductance.

**Figure 5.**
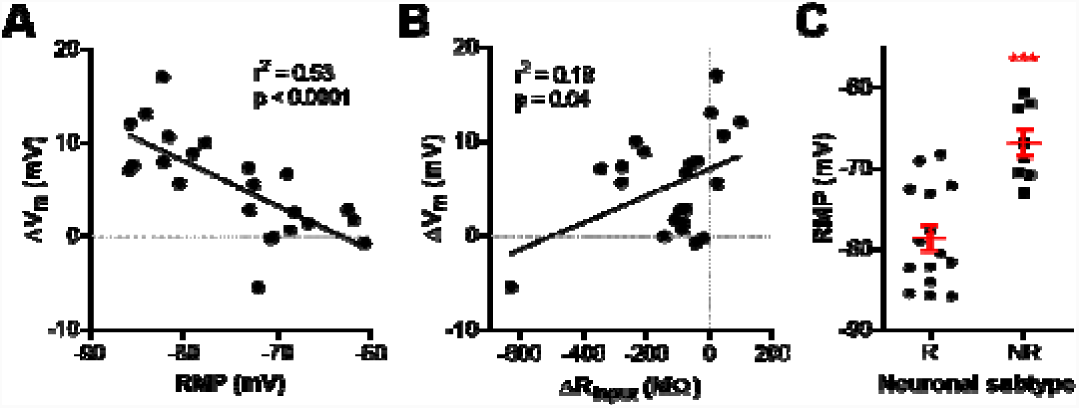
Low-glucose-induced depolarization of NTS neurons is a voltage-dependent mechanism. The graphs show the linear correlation between the change of membrane potential (ΔV_m_) induced by low glucose and (**A**) resting membrane potential (RMP) and (**B**) input resistance response (ΔR_input_) of neurons. **C,** comparison of the RMP between neurons responsive (R) and unresponsive (NR) to low glucose. ***p < 0.001.

In order to know if the decrease in glucose is being sensed by the recorded neuron itself, or signalizing by neighboring glia as previously suggested (McDougal et al., 2013) we added 3 mM of glucose in the recording pipette and measured the response of the neuron to low-glucose (n = 6, four animals). In this condition perfusion of low-glucose external solution did not change the RMP of NTS neurons (−1.0 ± 0.64 mV; from a mean of −76.3 ± 3.6 mV to −75.3 ± 3.2 mV; p = 0.2; **Figure 6A**), showing that the drop in external glucose is being sensed by the recorded neuron. The RMP of the neurons with 3 mM glucose was not significantly different from the RMP of NTS neurons in 5 mM glucose (p = 0.6; **Figure 6B**) showing that the presence of glucose in the pipette did not depolarize the neuron. In fact when we compared the RMP of the non-responsive neurons, which were more depolarized, with the RMP in 3 mM internal glucose, we found a significant difference (−66.8 ± 1.6 mV vs. −76.3 ± 3.6 mV, respectively; p = 0.02; **Figure 6C**) confirming that the internal glucose is not depolarizing the NTS neurons.

**Figure 6.**
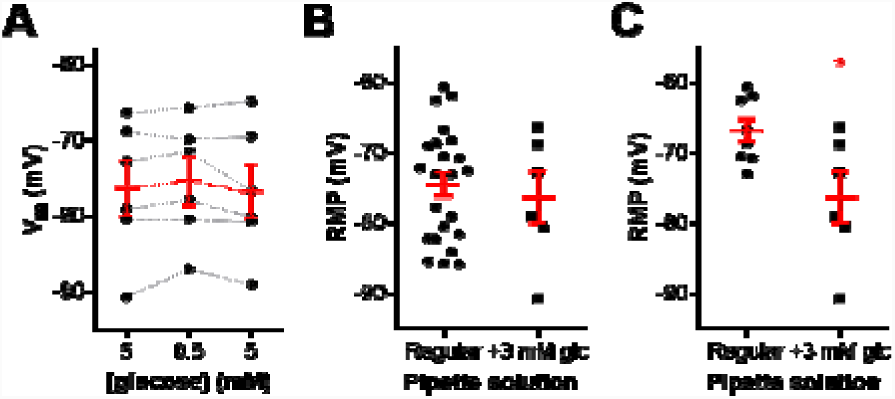
The presence of glucose in the pipette solution prevents the detection of low-external glucose by NTS neurons. **A,** summary of the low-glucose effect on the membrane potential (V_m_) of neurons recorded with 3 mM glucose-added pipette solution. **B,** comparison of the resting membrane potential (RMP) between neurons recorded with the regular pipette solution and the 3 mM glucose-added pipette solution. **C,** comparison of the RMP between unresponsive neurons recorded with the regular pipette solution and the 3 mM glucose-added pipette solution. Glc, glucose. *p < 0.05.

### Perfusion with low-glucose solution produces an inward current

We then investigated which ionic conductance is responsible for the depolarization induced by low glucose. Lamy et al., (2014) showed that in GABAergic neurons from mice NTS, low-glucose solution inhibits a potassium leak conductance, leading to a depolarization of the neuron. On the other hand, Balfour and Trapp (2007) reported in NTS neurons from rats an inward current activated by low glucose in half of the neurons, and an inhibition of a potassium conductance or a parallel inward current, suggestive of an inhibition of the current of the Na/K-ATPase. We performed current-voltage relationships voltage-clamp recordings in NTS neurons (n = 8, five animals), and observed a similar effect than observed by Balfour and Trapp, with low glucose aCSF perfusion increased membrane conductance from 3.1 ± 0.1 nS to 4.8 ± 0.1 nS producing a small inward current (−57.6 pA at −85 mV) with a reversion around –51 mV (**Figure 7A** and **7B**). Interestingly the reversal potential is around the membrane potential where the effect of low-glucose is not observed. Thus, we conclude that in NTS neurons from rats, the main mechanism of depolarization is the development of an inward current with a reversal potential more positive than the resting membrane potential. This is in accordance with the decrease in the input resistance observed after low glucose external solution in NTS neurons, and with the voltage-dependency of the depolarization.

**Figure 7.**
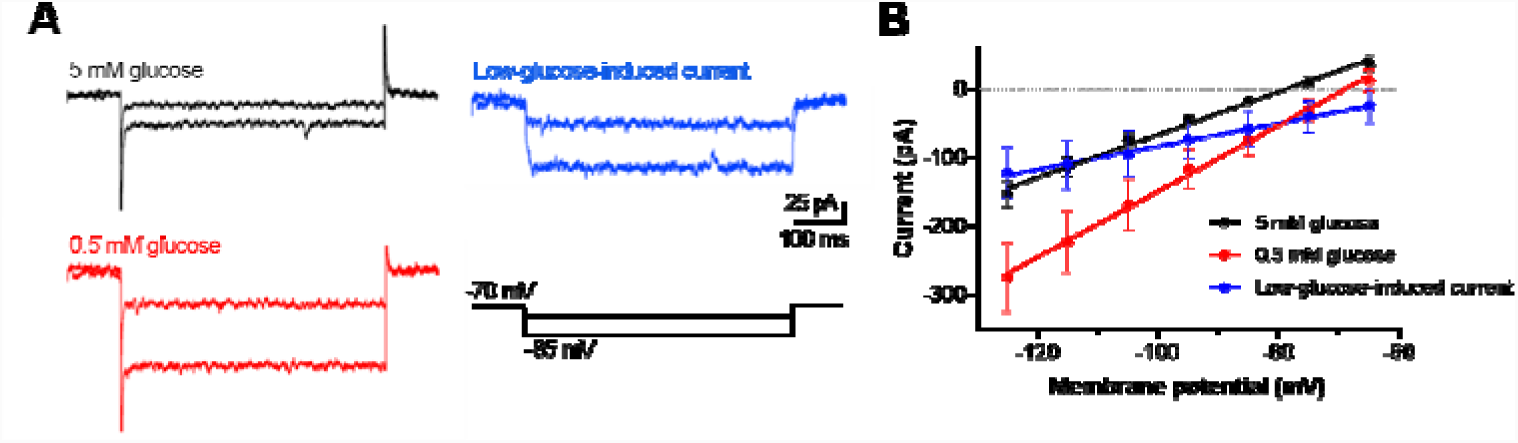
Low-glucose induced depolarization of NTS neurons is triggered by an inward current. **A,** representative time course recordings of currents in response to voltage steps obtained in basal (5 mM glucose; black) and low-glucose (0.5 mM; red) conditions, and the respective subtraction of currents (blue). **B,** linear regression of current-voltage relationship of neurons recorded in the conditions depicted in **A**.

### Depolarization by blocking K_ATP_ channels inhibits the effect of low-glucose aCSF

We showed that NTS neurons have K_ATP_ channels opened at rest and that these channels could strongly affect RMP if inhibited, as demonstrated by the depolarization triggered by their antagonist tolbutamide. Additionally, we found that perfusion with low glucose aCSF depolarized the RMP of these neurons, but this effect decreased with RMP depolarization, and the non-responsive neurons were more depolarize than the responsive neurons. Thus, two opposite signals, low-glucose and block of K_ATP_ channels produce a similar effect, the membrane depolarization. Because the depolarization induced by low external glucose is more prominent in more hyperpolarized neurons, in a situation where K_ATP_ channels were blocked, as in a high ATP/ADP ratio, the neuron could not sense the drop in external glucose.

In order to test this hypothesis, we first tested if a drop in the external glucose could further depolarize a neuron depolarized by tolbutamide. We first tested the responsiveness of the neuron to low glucose, washed it, and then applied tolbutamide. We found that all neurons responsive to low glucose were responsive to tolbutamide (**Figures 8A*i*** and **8A***ii*). Low glucose reversibly depolarized the RMP by 9.1 ± 2.2 mV (p = 0.01; n = 5), and application of tolbutamide was able to depolarize the RMP by 18.5 ± 3.0 mV (p = 0.004), showing that neurons responsive to low glucose have K_ATP_ channels opened at rest. However, in these neurons, application of low glucose aCSF in the presence of tolbutamide induced only a small, but significant, membrane depolarization (2.0 ± 0.3 mV; p = 0.002), which was smaller than the depolarizing response before tolbutamide application (p = 0.03; **Figure 8A*iii***). This result suggests that depolarization induced by blocking K_ATP_ channels blunts the neuron’s response to low external glucose.

**Figure 8.**
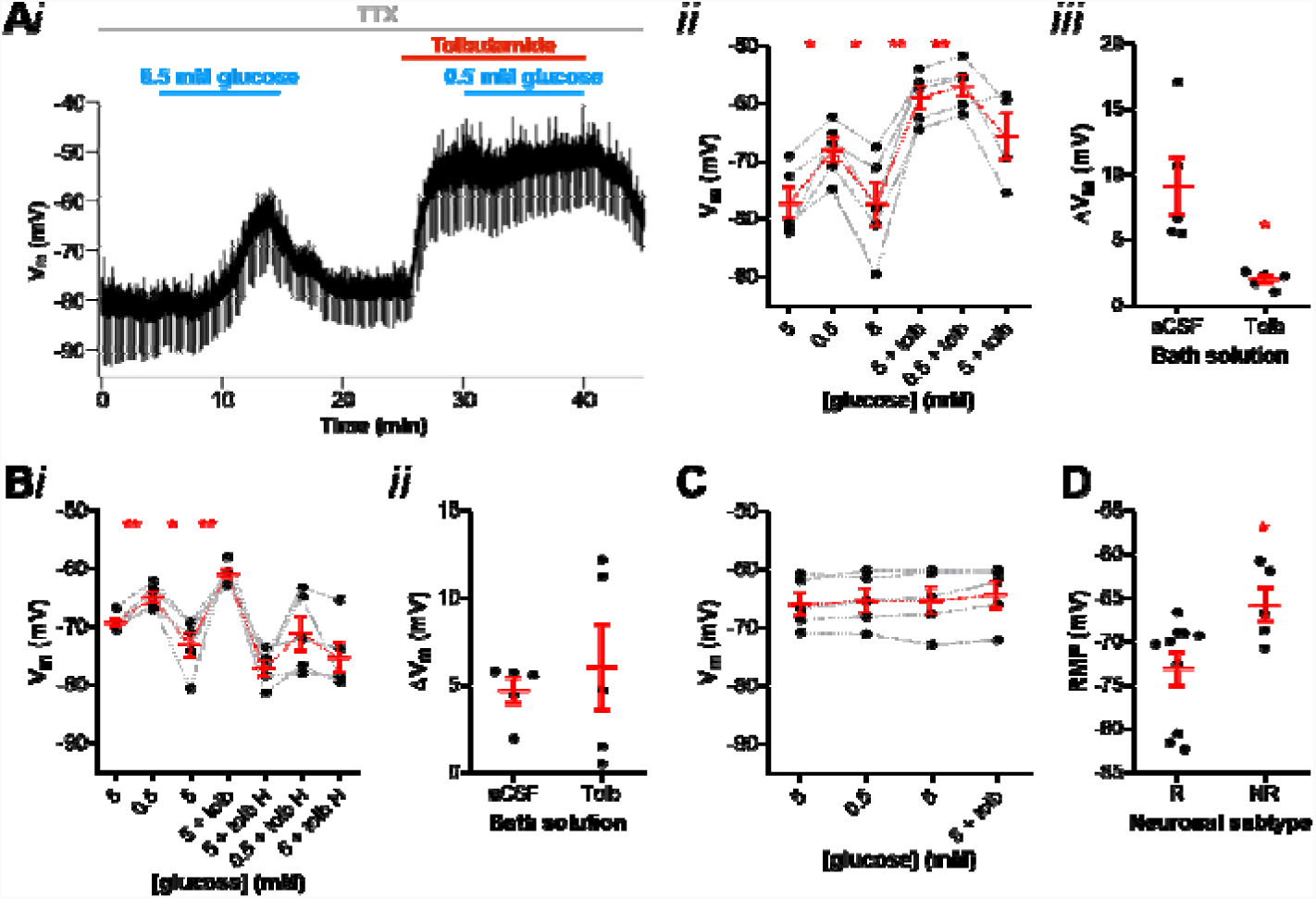
The occlusion of K_ATP_ channels suppresses the low-glucose induced depolarization of NTS neurons. **A,** application of tolbutamide blunts the depolarizing effect induced by a low-glucose challenge, as shown by a representative recording (*i*). Neurons responsive to low glucose are also depolarized by tolbutamide (*ii*), but this effect occludes the low-glucose sensing (*iii*). **B,** hyperpolarization of neurons responsive to low glucose (*i*) prevents the effect of tolbutamide in blunting the low-glucose sensing (*ii*). **C,** neurons unresponsive to low glucose do not also respond to tolbutamide application. **D,** comparison of the resting membrane potential (RMP) between neurons responsive (R) and unresponsive (NR) to both low glucose and tolbutamide. aCSF, artificial cerebrospinal fluid; H, hyperpolarization; Tolb, tolbutamide; TTX, tetrodotoxin; V_m_, membrane potential. *p < 0.05; **p < 0.01.

To test if this effect is only due to the membrane depolarization induced by tolbutamide we repeated this protocol, but hyperpolarized the neurons by injecting DC current after the application of tolbutamide, and tested the effect of low-glucose aCSF (n = 5; **Figure 8B*i***). Again, we found that the membrane potential is depolarized by both low glucose (4.4 ± 0.7 mV; p = 0.003) and tolbutamide (12.2 ± 2.8 mV; p = 0.001). Next, the injection of −30 to −50 pA of DC current hyperpolarized the membrane potential (from a mean of −60.5 ± 0.8 mV to −76.9 ± 1.3 mV), and in this condition, low-glucose aCSF was able to depolarize the membrane in more than 2 mV in 3 out of 5 NTS neurons, (average for all neurons: 6.0 ± 2.4 mV; p = 0.06;). However, this depolarization was similar to the observed in the absence of tolbutamide (p = 0.6; **Figure 8B*ii***) showing that tolbutamide is not able to occlude the effect of low glucose if the membrane is hyperpolarized to values similar to the RMP in most NTS neurons.

Interestingly, all neurons unresponsive to low glucose aCSF (0.6 ± 0.5 mV; p = 0.3; n = 5) were also not responsive to tolbutamide (1.0 ± 0.5 mV; p = 0.1; **Figure 8C**). These neurons also had more depolarized RMP in comparison to responsive neurons (−65.7 ± 1.9 mV vs. −73.1 ± 1.9 mV, respectively; p = 0.03; **Figure 8D**).

### Incubation with 10 mM glucose aCSF increases RMP and reduces the number of neurons responsive to low glucose and tolbutamide

Because most NTS neurons express open K_ATP_ channels at rest, which are able to depolarize the membrane when blocked, we asked if NTS neurons incubated with a higher glucose concentration could lead to more ATP production by the glycolytic/oxidative metabolism and blockage of the K_ATP_ channels and membrane depolarization. For this, we incubated the slices in aCSF solution containing twice the concentration of glucose (10 mM glucose). We found that NTS neurons incubated in 10 mM glucose aCSF (n = 17, eight animals) had more depolarized RMP than neurons incubated in 5 mM glucose aCSF (−69.0 ± 1.5 mV vs. −74.5 ± 1.7, respectively; p = 0.02; **Figure 9A**. This suggests that NTS neurons incubated in 10 mM glucose produce more ATP, which block K_ATP_ channels and depolarize the membrane. Because the response to low-glucose was blunted by membrane depolarization we hypothesized that NTS neurons incubated in 10 mM glucose would be less responsive to a low-glucose solution than neurons incubated in 5 mM glucose. In order to test this we sequentially applied a low-glucose solution, followed, after returning to 10 mM glucose, by application of tolbutamide. We found that only 3 out of 16 cells (19%) were reversible depolarized by perfusion of 0.5 mM glucose (11.2 ± 0.8 mV, from a mean of −74.0 ± 3.7 mV to −62.8 ± 3.3; p = 0.0008. **Figure 9B,C**). This depolarization was similar to what found in neurons incubated in 5 mM external glucose (p = 0.2). Interestingly, the time course of the depolarization was slower, taking on average 435 ± 39 ms to peak in contrast to 349 ± 19 ms in 5 mM glucose aCSF (p = 0.048). After returning to 10 mM glucose the RMP returned to a value similar to the original RMP (−72.7 ± 5.3 mV). Additionally, we sequentially applied tolbutamide which triggered a strong depolarization of 18.4 ± 1.6 mV (p = 0.007; **Figure 9B**), a similar value produced by tolbutamide in 5 mM glucose (p = 0.5). On the other hand, 13 neurons (81%) were considered unresponsive to low glucose (**Figure 9C**; 1.4 ± 0.5 mV, from a mean of −67.5 ± 1.4 mV to −66.2 ± 1.4 mV), and had more depolarized RMP compared to the responsive cells (responsive neurons: −77.7 ± 0.5 mV; unresponsive neurons: −66.05 ± 2.2 mV; p = 0.01; **Figure 9D**). Moreover, tolbutamide was applied in four of the unresponsive cells and did not induce significant membrane potential change (0.8 ± 0.3 mV; p = 0.08; **Figure 9E**). Like in neurons incubated with 5 mM glucose, we found a positive correlation between the effect of low glucose and RMP (r^2^ = 0.26, p = 0.04; **Figure 9F**).

**Figure 9.**
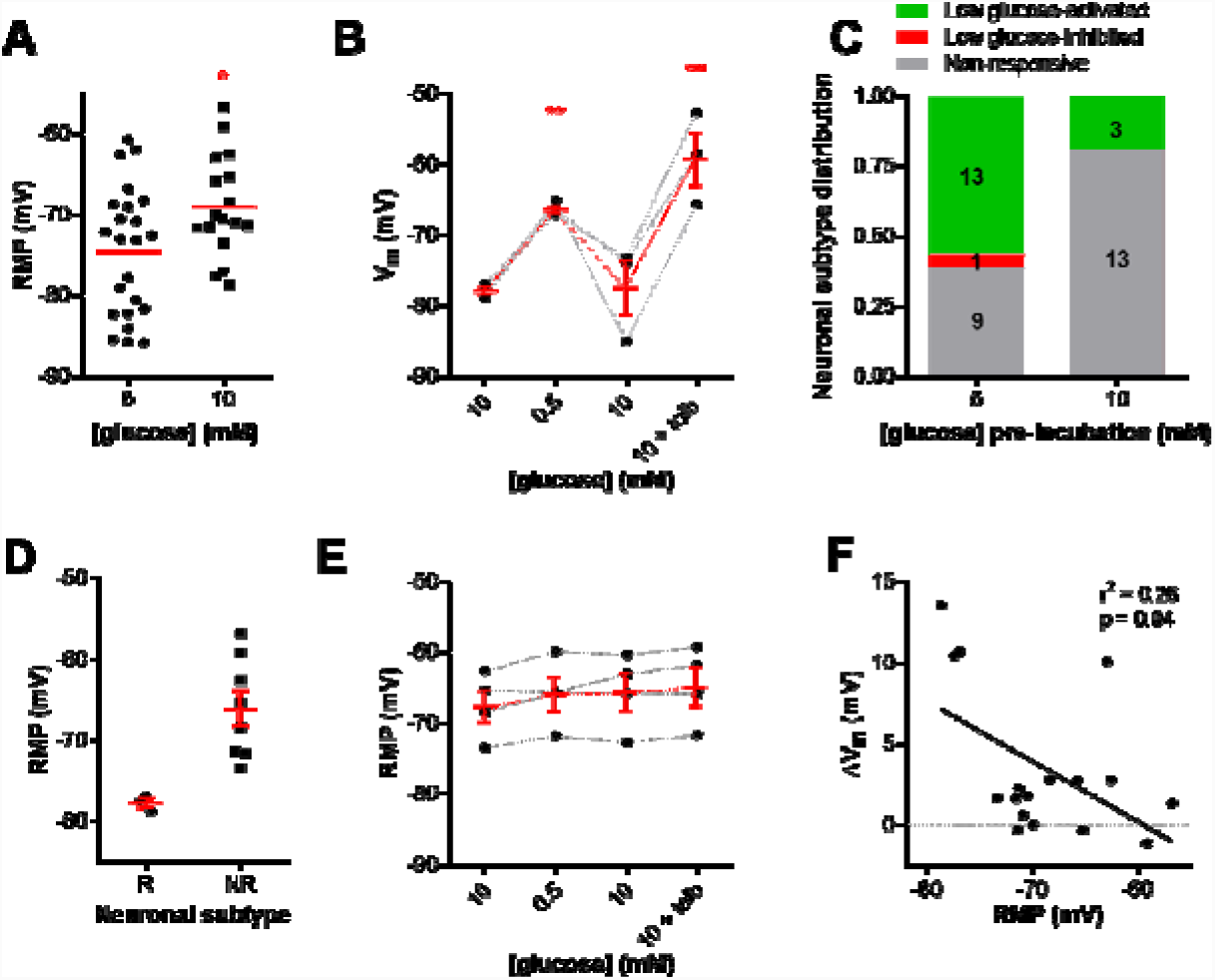
Incubation in high glucose (10 mM) decreases the number of NTS neurons sensitive to low glucose. **A,** comparison of the resting membrane potential (RMP) between neurons incubated in 5 mM and 10 mM glucose. **B,** neurons incubated in high glucose that are responsive to low glucose are depolarized by tolbutamide application. **C,** comparison of the distribution of neurons pre-incubated in different glucose solutions according to their response to low glucose. **D,** comparison of the RMP of neurons incubated in high glucose that responsive (R) or unresponsive (NR) to a low-glucose challenge. **E,** neurons incubated in high glucose that are unresponsive to low glucose do not also respond to tolbutamide application. **F,** linear correlation between the change of membrane potential (V_m_) and the RMP of neurons incubated incubated in high glucose. Tolb, tolbutamide. *p < 0.05; **p < 0.01.

The lack of effect of both low glucose and tolbutamide on the RMP of NTS neurons suggest that metabolic ATP is blocking K_ATP_ in these neurons when incubated in 10 mM glucose, leading to the membrane depolarization and inhibition of the response to low-glucose. To test this we hyperpolarized the depolarized neurons (from −71.3 ± 3.5 mV to 85.1 ± 2.9 mV; n = 5) and applied sequentially low glucose and tolbutamide (**Figure 10A**). Surprisingly, in hyperpolarized neurons, the response to low glucose was existent but reduced when compared to regular responsive neurons (3.2 ± 1.1 mV vs. 11.2 ± 0.8 mV, respectively; p = 0.0004; **Figure 10B**), suggesting a diminished response to low glucose in neurons incubated in 10 mM glucose. On the other hand, these neurons continued to be non-responsive to tolbutamide (−2.4 ± 1.1 mV; from −83.0 ± 1.2 mV to −80.6 ±1.5 mV; p = 0.1; **Figure 10A,C**). Although some individual neurons presented a smaller response, they were much smaller than in 5 mM glucose (p = 0.03; **Figure 10C**). These results are in accordance to the hypothesis that metabolic ATP is blocking the K_ATP_ channels.

**Figure 10.**
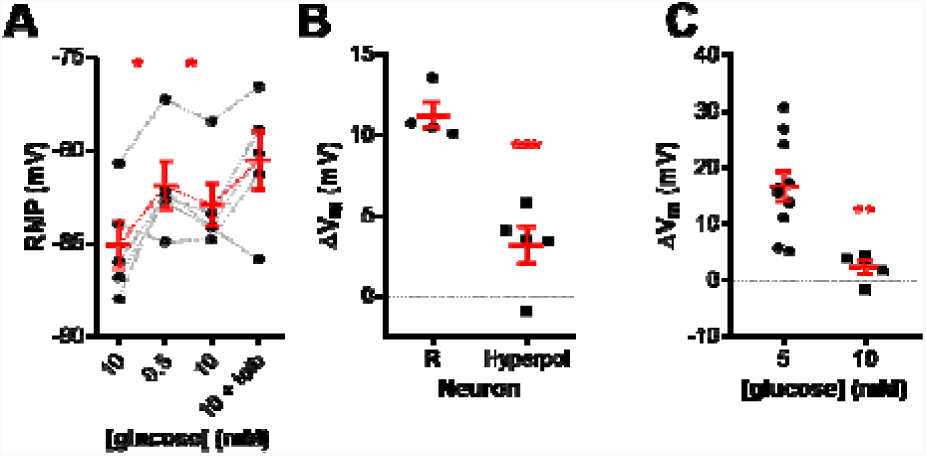
Hyperpolarization of NTS neurons incubated in high glucose (10 mM) does not rescue the response to a low-glucose challenge and tolbutamide application. **A,** neurons incubated in high glucose that had more positive resting membrane potential (RMP) were hyperpolarized and challenged to a low glucose solution and tolbutamide application, respectively. **B,** comparison of the membrane potential change (ΔV_m_) induced by low glucose between high-glucose-incubated responsive neurons (R) and the hyperpolarized neurons (Hyperpol) shown in **A**. **C,** comparison of the ΔV_m_ induced by tolbutamide between neurons pre-incubated in 5 mM and 10 mM glucose. Tolb, tolbutamide. *p < 0.05.

We conclude that incubating NTS neurons in a high glucose aCSF decreases the number of low glucose-responsive neurons. Although, this effect was not caused by the more depolarized membrane potential of these neurons, these neurons when hyperpolarized responded more weakly to low glucose. On the other hand, the more depolarized RMP suggest that the higher extracellular glucose can increase metabolic ATP, which blocks K_ATP_ channels and depolarize neuronal membrane.

### The depolarization induced by low glucose does not sustain for prolonged periods

Because of the substantial effect of K_ATP_ channels modulating the RMP of NTS neurons and the evidences above suggesting that the metabolic ATP can control RMP by modulating K_ATP_ channels, we asked if in a situation of prolonged exposure to a low glucose external solution, where metabolic ATP can be reduced, the membrane depolarization could be sustained.

For this, we recorded the membrane potential after perfusion of 0.5 mM glucose for longer than the 10-minute period we used in the previous recordings (n = 4). We observed in three neurons that the depolarization caused by perfusion with 0.5 mM glucose starts to revert around 1033 ± 101 seconds. In two of these neurons we applied tolbutamide and it depolarized the RMP to values similar to at the beginning of low glucose aCSF (**Figure 11A*i*** and **11A***ii*). Interestingly, in one non-responsive neuron, the RMP hyperpolarized after 1300 seconds of exposure to 0.5 mM glucose, and addition of tolbutamide reverted the hyperpolarization (**Figure 11B**), suggesting that this hyperpolarization was caused at least partially by the opening of K_ATP_ channels. To investigate this further we applied tolbutamide during the low-glucose depolarization (n = 3). In these three neurons, tolbutamide avoided the hyperpolarization caused by prolonged low glucose exposure (**Figure 11C***i* and **C***ii*). We conclude that the effect of low-external glucose in depolarizing the membrane of NTS neurons is short living and is probably reverted by depletion of intracellular ATP and opening of K_ATP_ channels.

**Figure 11.**
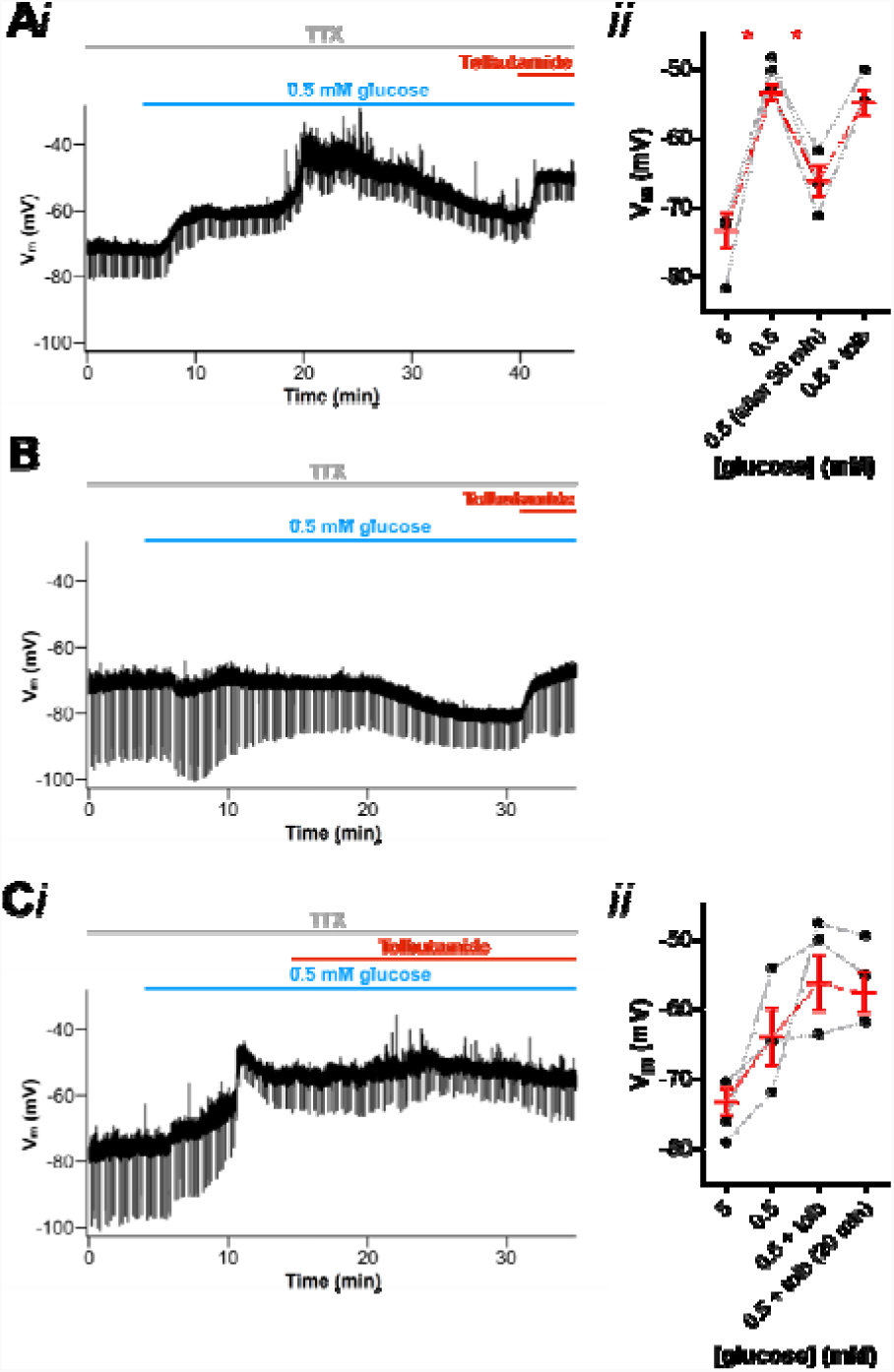
Low-glucose induced depolarization of NTS neurons is short living and reverted by the opening of K_ATP_ channels. **A,** subset of neurons responsive to low glucose is hyperpolarized after a long period exposed to it, as shown by a representative recording (*i*). Note that tolbutamide reverts the hyperpolarizing effect induced by low glucose. **A***ii*, summary of the effect triggered by low glucose and tolbutamide on the membrane potential (V_m_) of neurons. **B,** a non-responsive neuron to a low-glucose challenge is hyperpolarized after more than 20 min exposed to it. Note that tolbutamide also reverts the hyperpolarizing effect. **C,** application of tolbutamide suppresses the hyperpolarization induced by a long period of low-glucose perfusion, as shown by a representative recording (*i*). **C***ii*, summary of the effect of low glucose and tolbutamide on V_m_ of neurons. Tolb, tolbutamide; TTX, tetrodotoxin. *p < 0.05.

## Discussion

Glucose is the primary energy source for brain metabolism and survival (Mergenthaler *et al.*, 2013). Due to high levels of energy expenditure for neuronal activity and low content of brain glycogen, the human brain consumes up to 20% of the glucose-derived energy under physiological conditions (Magistretti & Allaman, 2015). Brain hypoglycemia, a condition of limited energy availability, can cause neuronal death and may lead to cognitive impairments and conscience loss (Cryer, 2007). Therefore, several peripheral and central components act on energy homeostasis regulation in order to maintain adequate levels of circulating glucose (Marty *et al.*, 2007; Verberne *et al.*, 2014).

Recent evidences have demonstrated that glucose-sensing neurons located in the brainstem nucleus of the tractus solitarius (NTS) can sense glucose levels in the extracellular milieu, using mechanisms which could involve or not K_ATP_ channels (Balfour *et al.*, 2006; Lamy *et al.*, 2014; Boychuk *et al.*, 2015; Halmos *et al.*, 2015; Roberts *et al.*, 2017). Here we found that NTS neurons express K_ATP_ channels as they all responded to diazoxide. In 5 mM glucose, most of these channels are open, since they depolarized in response to tolbutamide. Thus, in normoglycemic conditions, NTS neurons have open K_ATP_ channels which could be modulated by metabolism. In fact, when we incubated the slices in 10 mM glucose, the neurons were more depolarized and unresponsive (or much less responsive) to tolbutamide. These effects were not observed when the internal electrode solution contained 3 mM glucose, showing the drop in glucose was sensed by the neuron itself, and not by glia or other neurons in the circuit. These data show that NTS neurons can sense extracellular glucose and modulate their membrane potential using K_ATP_ channels.

Our experiments were performed in whole-cell patch-clamp using 4 mM Mg-ATP in the internal solution. Balfour *et al.*, 2006 showed that NTS neurons are more responsible to metabolic ATP than the ATP provded by the electrode. Similar results were described by Muller et al., (2002) in dorsal vagal neurons, and hypothalamic neurons (Song et al., 2000). Although it is not clear the reason of this effect it is probaly could be due to the compartimentalization of K_ATP_ channels in special membrane domains (Garg et al., 2009) which could be closer to the mitochondria and glycolitic enzymes which have been shown to concentrate in regions of high metabolic demand and glucose transport actvivity (Zechin etal., 2015, Agrawal et al., 2018). Additionally, this concentration of ATP is more than the micromolar concentrations needed to block isolated K_ATP_ channels (Inagaki et al., 1996), but the effect of toulbutamide show that there are K_ATP_ channels open even with 4 mM internal ATP. This can be explained by knowing that the K_ATP_ channels are sensitive to the ADP:ATP ratio, because they are activated by MgADP (Nichols et al., 1996; Shying et al., 1997) and their affinity to ATP is greatly reduced by phosphatidylinositol 4,5-bis-phosphate (PIP2)(Baukrowitz et al., 1998; Shyng et al., 2000). Thus, our whole cell recordings very likely mostly reflects the physiological responses of NTS neurons to metabolic ATP derived from external glucose.

In the current investigation, we showed that low glucose (0.5 mM) induces a voltage-dependent depolarization in most NTS neurons of rats. Additionally, we showed that the RMP in NTS neurons incubated in 10 mM glucose aCSF is more depolarized and less sensitive to tolbutamide, even when hyperpolarized, suggesting they are depolarized by blocking of K_ATP_ channels by metabolic ATP. Therefore, we expected that the depolarization of the RMP induced by the incubation of slices in high glucose could lead to an increase in the number of neurons unresponsive to a low-glucose challenge, since our findings demonstrated a voltage-dependent depolarization by low glucose (i.e., the more negative the RMP, the greater the membrane potential response amplitude), and this effect was indeed observed. Neurons unresponsive to low glucose accounted for 35% when incubated in normal aCSF, but ∼70% when incubated in high-glucose aCSF. Interestingly, Balfour *et al*. (2006) reported that ∼80% of NTS/DMX neurons of rats were unresponsive to glucose removal; however these authors performed the electrophysiological recordings using a control aCSF with 10 mM glucose. These authors also showed the expression of the K_ATP_ channel subunit SUR1 in cells that did not respond to low glucose, what suggest that in these neurons K_ATP_ channels are present but could be saturated by high glucose-derived ATP levels.

Low extracellular glucose induce K_ATP_ channels to open following a decrease in intracellular ATP levels (Hibino *et al.*, 2010), but we observed a hyperpolarization response to low glucose in only two of 60 NTS neurons incubated in 5 mM glucose aCSF. Other groups have reported that the sensitivity to glucose is not observed in all NTS neurons expressing K_ATP_ channel subunits (Dallaporta *et al.*, 2000; Balfour *et al.*, 2006). Lamy *et al*. (2014) reported an increase in membrane input resistance by the closure of leak potassium channels in GLUT2-expressing GABAergic neurons activated by low glucose in mice. This could account for the response we observed, but again, the low glucose-sensitive current observed in this report was a linear non-rectifying current, which is not compatible to the voltage-dependency we observed, and we observed mainly a decrease in input resistance, suggesting the opening of an inward conductance. Additionally, our findings demonstrate that most NTS neurons exhibit a voltage-dependent depolarization and a decrease in input resistance in response to a low-glucose solution, which is suggestive of the opening of a cationic conductance. We found that low glucose induces an inward cationic current with a reversion around −60 mV, which could explain the voltage-dependence of the depolarization induced by low glucose. Interestingly, Balfour *et al*. (2007) showed the opening of an inwardly rectifying current in some neurons of the NTS in response to glucose removal, what could contribute to the depolarizing response seen in our recordings. The opening of HCN channels which mediated the inwardly rectifying cationic h current could explain our results, but the activation of the low-glucose induced current was very fast, not compatible with the slow activation of the HCN channels. Additionally, we performed experiments using the antagonist of these channels ZD7288 (10 µM), but the results were inconclusive because this drug produces a progressive depolarization of the membrane masking any effect of low-glucose (not shown). In the ventromedial hypothalamus, low glucose lead to the closure of chloride channels and membrane depolarization in some neurons (Routh *et al.*, 2014), but the reversal potential of the low glucose induce current observed by us does not suggest a chloride conductance. Other ionic mechanisms than ion channels can contribute to depolarize neurons under low-glucose availability. The Na^+^/K^+^-ATPase pumps are important generators of electrochemical gradients in cells. High-glucose levels stimulate the Na^+^/K^+^-ATPase pump and triggers membrane hyperpolarization in glucose-excited neurons in the lateral hypothalamus (Oomura *et al.*, 1974), and brain hypoglycemia reduces the activity of the Na^+^/K^+^-ATPase pump (Lees, 1991) and may lead to depolarization of neurons (Balfour & Trapp, 2007). However, this mechanism is not compatible to the voltage-dependency and the decrease in the input resistance we observed. Nevertheless, the reduction of the activity of the Na^+^/K^+^-ATPase pump could constitute an additional component altering the RMP of NTS neurons in response to reduced external glucose concentration.

Since both K_ATP_ blockage and low glucose either produce membrane depolarization, but low glucose is ineffective to depolarize neurons when K_ATP_ are fully blocked, we believe that the depolarization induced by low external glucose might represent a form of homeostatic regulation of the RMP in order to avoid an excessive hyperpolarization by unblocking K_ATP_ channels under a hypoglycemia episode, which could be essential for the proper functioning of the vital autonomic functions controlled by the NTS. However when we left the neurons for more than 10-20 minutes in low external glucose, we started to observe a hyperpolarization which was prevented by tolbutamide, showing that the low-glucose depolarization does not last for prolonged periods and is not effective in a long-term period, and the membrane starts to hyperpolarize in response to the decreased metabolic ATP.

Our results clear show a mechanism of membrane depolarization driven by a rapid reduction in external glucose that could be potentially relevant for neurometabolic responses to hypoglycemia. However, when neurons are maintained in a hyperglycemic solution (similar to high post-prandial circulating glucose levels and diabetic conditions), the number of NTS neurons responsive to low glucose decreases, and the latency for peak response increases. Because NTS neurons are involved in the counterregulatory mechanisms to hypoglycemia, which includes a decrease in pancreatic insulin secretion, an increase in pancreatic glucagon secretion, and increase in adrenomedullary epinephrine secretion (Cryer et al., 2005), the detection of a rapid drop in glicemia by these neurons in prolonged hyperglycemic situations might be blunted. Sudden drops in glycemia is a common occurrence in patients with type 1 and advanced Type II diabetes, caused by insulin interventions to control glycemia (Mergenthaler *et al.*, 2013). In these patients, a failed counterregulatory response to hypoglycemia is a common problem, a condition named hypoglycemia-associated autonomic failure (Cryer, 2005). Our data suggests that the establishment of more depolarized RMP of NTS neurons by the closure of K_ATP_ channels due to increased intracellular ATP levels could be an additional mechanism contributing to reduced brain sensitivity to a low-glucose challenge during hyperglycemic conditions. More studies are needed to support this interesting hypothesis.

## Acknowledgements

We thank the technical support of Mr. J. Fernando Aguiar, and Drs. Daniela Accorsi-Mendonça, Luiz C. Navegantes and Luiz G. Branco for their criticisms and suggestions. Research supported by FAPESP grant (2016/01607-4). CBM is a CNPq PhD scholarship recipient. RML is a CNPq research fellow.

## References

Agrawal A, Pekkurnaz G, Koslover EF (2018). Spatial control of neuronal metabolism through glucose-mediated mitochondrial transport regulation. Elife. 18;7.

Accorsi-Mendonça D, Castania JA, Bonagamba LG, Machado BH, Leão RM (2011). Synaptic profile of nucleus tractus solitarius neurons involved with the peripheral chemoreflex pathways. Neuroscience. 197:107–20.

Andrew, S. F., Dinh, T. T., and Ritter, S. (2007). Localized glucoprivation of hindbrain sites elicits corticosterone and glucagon secretion. Am. J. Physiol. Regul. Integr. Comp. Physiol. 292, R1792–1798.

Ashcroft FM & Rorsman P. (2013). K(ATP) channels and islet hormone secretion: new insights and controversies. Nat Rev Endocrinol 9, 660–669.

Balfour RH, Hansen AM & Trapp S. (2006). Neuronal responses to transient hypoglycaemia in the dorsal vagal complex of the rat brainstem. J Physiol 570, 469–484.

Balfour RH & Trapp S. (2007). Ionic currents underlying the response of rat dorsal vagal neurones to hypoglycaemia and chemical anoxia. J Physiol 579, 691–702.

Bohland M, Matveyenko AV, Saberi M, Khan AM, Watts AG & Donovan CM. (2014). Activation of hindbrain neurons is mediated by portal-mesenteric vein glucosensors during slow-onset hypoglycemia. Diabetes 63, 2866–2875.

Boychuk CR, Gyarmati P, Xu H & Smith BN. (2015). Glucose sensing by GABAergic neurons in the mouse nucleus tractus solitarii. J Neurophysiol 114, 999–1007.

Baukrowitz T., Schulte U., Oliver D., Herlitze S., Krauter T., Tucker S. J., Ruppersberg J.P., Fakler B. (1998). PIP2 and PIP as determinants for ATP inhibition of KATP channels. Science 282, 1141–1144.

Cryer PE. (2005). Mechanisms of hypoglycemia-associated autonomic failure and its component syndromes in diabetes. Diabetes 54, 3592–3601.

Cryer PE. (2007). Hypoglycemia, functional brain failure, and brain death. J Clin Invest 117, 868–870.

Dallaporta M, Perrin J & Orsini JC. (2000). Involvement of adenosine triphosphate-sensitive K+ channels in glucose-sensing in the rat solitary tract nucleus. Neurosci Lett 278, 77–80.

Donovan CM & Watts AG. (2014). Peripheral and central glucose sensing in hypoglycemic detection. Physiology (Bethesda) 29, 314–324.

Gao L, Ortega-Sáenz P, García-Fernández M, González-Rodríguez P, Caballero-Eraso C & López-Barneo J. (2014). Glucose sensing by carotid body glomus cells: potential implications in disease. Front Physiol 5, 398.

Garg V, Jiao J, Hu K. (2009) Regulation of ATP-sensitive K+ channels by caveolin-enriched microdomains in cardiac myocytes. Cardiovasc Res. 82:51–8.

Gross, P. M., Wall, K. M., Pang, J. J., Shaver, S. W., and Wainman, D. S. (1990). Microvascular specializations promoting rapid interstitial solute dispersion in nucleus tractus solitarius. Am. J. Physiol. 259, R1131–R1138

Halmos KC, Gyarmati P, Xu H, Maimaiti S, Jancsó G, Benedek G & Smith BN. (2015). Molecular and functional changes in glucokinase expression in the brainstem dorsal vagal complex in a murine model of type 1 diabetes. Neuroscience 306, 115–122.

Hermann GE, Viard E & Rogers RC. (2014). Hindbrain glucoprivation effects on gastric vagal reflex circuits and gastric motility in the rat are suppressed by the astrocyte inhibitor fluorocitrate. J Neurosci 34, 10488–10496.

Herzog RI, Jiang L, Herman P, Zhao C, Sanganahalli BG, Mason GF, Hyder F, Rothman DL, Sherwin RS & Behar KL. (2013). Lactate preserves neuronal metabolism and function following antecedent recurrent hypoglycemia. J Clin Invest 123, 1988–1998.

Hibino H, Inanobe A, Furutani K, Murakami S, Findlay I & Kurachi Y. (2010). Inwardly rectifying potassium channels: their structure, function, and physiological roles. Physiol Rev 90, 291–366.

Inagaki N, Gonoi T, Clement JP, Wang CZ, Aguilar-Bryan L, Bryan J, Seino S. (1996). A family of sulfonylurea receptors determines the pharmacological properties of ATP-sensitive K+ channels. Neuron. 16:1011–7

Lam CK, Chari M, Su BB, Cheung GW, Kokorovic A, Yang CS, Wang PY, Lai TY & Lam TK. (2010). Activation of N-methyl-D-aspartate (NMDA) receptors in the dorsal vagal complex lowers glucose production. J Biol Chem 285, 21913–21921.

Lamy CM, Sanno H, Labouèbe G, Picard A, Magnan C, Chatton JY & Thorens B. (2014). Hypoglycemia-activated GLUT2 neurons of the nucleus tractus solitarius stimulate vagal activity and glucagon secretion. Cell Metab 19, 527–538.

Lees GJ. (1991). Inhibition of sodium-potassium-ATPase: a potentially ubiquitous mechanism contributing to central nervous system neuropathology. Brain Res Brain Res Rev 16, 283–300.

Magistretti PJ & Allaman I. (2015). A cellular perspective on brain energy metabolism and functional imaging. Neuron 86, 883–901.

Marty N, Dallaporta M & Thorens B. (2007). Brain glucose sensing, counterregulation, and energy homeostasis. Physiology (Bethesda) 22, 241–251.

McDougal DH, Hermann GE & Rogers RC. (2013). Astrocytes in the nucleus of the solitary tract are activated by low glucose or glucoprivation: evidence for glial involvement in glucose homeostasis. Front Neurosci 7, 249.

Mergenthaler P, Lindauer U, Dienel GA & Meisel A. (2013). Sugar for the brain: the role of glucose in physiological and pathological brain function. Trends Neurosci 36, 587–597.

Müller M, Brockhaus J, Ballanyi K. (2002) ATP-independent anoxic activation of ATP-sensitive K+ channels in dorsal vagal neurons of juvenile mice in situ. Neuroscience. 109:313–28.

Nichols, C.G., Shyng S.-L., Nestorowicz A., Glaser B., Clement IV J., Gonzalez G., Aguilar-Bryan L.,. Permutt A.M, Bryan J.P. (1996) Adenosine diphosphate as an intracellular regulator of insulin secretion. Science. 272:1785–1787

Nichols CG. (2006). KATP channels as molecular sensors of cellular metabolism. Nature 440, 470–476.

Nortley R & Attwell D. (2017). Control of brain energy supply by astrocytes. Curr Opin Neurobiol 47, 80–85.

Oliveira-Maia AJ, Roberts CD, Simon SA & Nicolelis MA. (2011). Gustatory and reward brain circuits in the control of food intake. Adv Tech Stand Neurosurg 36, 31–59.

Oomura Y, Ooyama H, Sugimori M, Nakamura T & Yamada Y. (1974). Glucose inhibition of the glucose-sensitive neurone in the rat lateral hypothalamus. Nature 247, 284–286.

Ritter S, Dinh TT & Zhang Y. (2000). Localization of hindbrain glucoreceptive sites controlling food intake and blood glucose. Brain Res 856, 37–47.

Roberts BL, Zhu M, Zhao H, Dillon C & Appleyard SM. (2017). High glucose increases action potential firing of catecholamine neurons in the nucleus of the solitary tract by increasing spontaneous glutamate inputs. Am J Physiol Regul Integr Comp Physiol 313, R229–R239.

Routh VH, Hao L, Santiago AM, Sheng Z & Zhou C. (2014). Hypothalamic glucose sensing: making ends meet. Front Syst Neurosci 8, 236.

Seaquist ER, Damberg GS, Tkac I, Gruetter R. (2011) The effect of insulin on in vivo cerebral glucose concentrations and rates of glucose transport/metabolism in humans. Diabetes, 50:2203–9

Shyng S Ferrigni T, Nichols CG. (1997) Regulation of KATP channel activity by diazoxide and MgADP. Distinct functions of the two nucleotide binding folds of the sulfonylurea receptor. J Gen Physiol. 110:643–54.

Shyng SL, Barbieri A, Gumusboga A, Cukras C, Pike L, Davis JN, Stahl PD, Nichols CG. (2000) Modulation of nucleotide sensitivity of ATP-sensitive potassium channels by phosphatidylinositol-4-phosphate 5-kinase. Proc Natl Acad Sci U S A. 97:937–41.

Song Z, Levin BE, McArdle JJ, Bakhos N, Routh VH. (2001) Convergence of pre- and postsynaptic influences on glucosensing neurons in the ventromedial hypothalamic nucleus. Diabetes. 50:2673–81.

Song Z & Routh VH. (2006). Recurrent hypoglycemia reduces the glucose sensitivity of glucose-inhibited neurons in the ventromedial hypothalamus nucleus. Am J Physiol Regul Integr Comp Physiol 291, R1283–1287.

Thorens B. (2012). Sensing of glucose in the brain. Handb Exp Pharmacol, 277–294.

Verberne AJ, Sabetghadam A & Korim WS. (2014). Neural pathways that control the glucose counterregulatory response. Front Neurosci 8, 38.

Zecchin A, Stapor PC, Goveia J, Carmeliet P. (2015) Metabolic pathway compartmentalization: an underappreciated opportunity? Curr Opin Biotechnol. 34:73–81.

Zhao S, Kanoski SE, Yan J, Grill HJ & Hayes MR. (2012). Hindbrain leptin and glucagon-like-peptide-1 receptor signaling interact to suppress food intake in an additive manner. Int J Obes (Lond) 36, 1522–1528.

